# OxLDL-targeted Chimeric Antigen Receptor T Regulatory Cells Reduce Atherosclerotic Plaque Development

**DOI:** 10.1101/2025.01.24.634830

**Authors:** Robert D. Schwab, Seok-Jae Albert Hong, Xin Bi, David A. Degaramo, Aisha Faruqi, Shawna K. Brookens, John T. Keane, Fang Liu, Kiran Musunuru, Daniel J. Rader, Avery D. Posey

## Abstract

Cardiovascular disease caused by atherosclerosis is responsible for 18 million deaths annually, highlighting a significant need for new medical therapies, especially for patients ineligible for surgical interventions. Atherosclerosis is driven by the accumulation of low-density lipoprotein (LDL) and the formation of foam cells, accompanied by oxidative stress and the deposition of oxidized LDL (OxLDL), a pro-inflammatory molecule. Lowering LDL is the mainstay of current medical treatment in addition to blood pressure control and lifestyle changes, but to date, specifically targeting the inflammatory pathways contributing to plaque development without significant systemic side effects has not been feasible. Over the past decade, chimeric antigen receptor (CAR) T cells have treated cancer and restored immune imbalance in autoimmune diseases in patients and resolved cardiac fibrosis in preclinical models. Using an inducible T regulatory cell (Treg) platform, we created an anti-OxLDL-specific CAR Treg therapy that exerts cell- and cytokine-mediated immunosuppression to reduce macrophage-foam cell formation in vitro. Murine anti-OxLDL CAR Tregs inhibited 80% of atherosclerotic plaque formation in immunocompetent mouse models of hyperlipidemia and atherosclerosis. These studies illustrate the potential of anti-OxLDL CAR Tregs to mitigate the inflammation and plaque deposition associated with OxLDL, potentially offering a new therapeutic option for atherosclerosis.

## Introduction

Atherosclerosis is a narrowing and hardening of arteries that can eventually lead to thrombotic events, causing inefficient oxygen penetration into essential tissues^1^. This hardening, plaque formation, and eventual rupture are due to progressive endothelial dysfunction from cholesterol deposition, resulting in inflammation and immune deregulation^2^. Oxidation of LDL by reactive oxygen species or nearby enzymes in the arterial wall and subsequent accumulation of OxLDL by macrophages is implicated as one of the initiating factors of atherosclerosis^3,4^. OxLDL is a primary pro-inflammatory signal that activates the endothelium, induces cytokine and chemokine release, and recruits monocytes, macrophages, and T-helper 1 cells^5,6^. OxLDL is normally catabolized to free cholesterol and free fatty acids in the late endosomes of macrophages and then secreted; however, in atherosclerosis, high concentrations of OxLDL lead to the accumulation of intracellular cholesterol crystals and lipid droplets, which transforms the macrophages into foam cells^7^. These foam cells and other antigen-presenting cells recruit and activate T and B lymphocytes, producing pro-inflammatory cytokines and atherogenic IgG immunoglobulins^8^. Eventually, foam cells undergo apoptosis and create a necrotic core, which promotes increased inflammation, metalloproteinase release, and smooth muscle injury^9^. In time, a fibrous cap develops on the arterial wall, leading to plaque deposition and rupture^10^.

Reducing LDL cholesterol is proven to reduce the progression of atherosclerosis and the risk of atherosclerotic cardiovascular events^11^. However, in patients with known atherosclerosis, the risk of CV events remains high even in the setting of well-controlled LDL cholesterol levels, indicating that new treatment paradigms are needed^12^. Inflammation is a major contributor to the atherosclerotic process and acute CV events, which are primarily driven by response to OxLDL^13^. OxLDL is elevated under chronic inflammatory conditions such as psoriasis, rheumatoid arthritis, HIV, and systemic lupus erythematosus^14–17^, and a recent meta-analysis demonstrated that elevated OxLDL levels in chronically inflamed conditions were significantly associated with higher rates of cardiovascular disease^18^.

Several studies have also shown the utility of OxLDL as a predictive biomarker^19^ and therapeutic target. For instance, targeted blockade of OxLDL in hypercholesterolemic mice, either through antibody infusion or genetic engineering for constitutive hepatocyte-specific antibody secretion, significantly reduced atherosclerotic lesion size and plaque inflammation^20,21^. Additionally, an anti-OxLDL-IL-10 immunoconjugate demonstrated reduced vascular inflammation in proatherogenic mice^22^. In a randomized, double-blind, placebo-controlled phase 2a trial in subjects with psoriasis treated with an Orticumab, a fully human monoclonal antibody against a specific OxLDL epitope, there was a reduction in coronary inflammation^23^. In patient samples, titers of anti-OxLDL IgG antibodies positively correlate with atherosclerosis and are pro-atherogenic^24^. In contrast, anti-OxLDL IgM antibodies are inversely associated with carotid intima-media thickness and may have atheroprotective effects^25^. In a pro-atherogenic mouse model, transferring IgM-secreting B1a lymphocytes reduced atherosclerotic lesions and decreased OxLDL and apoptotic cells^26^. Similarly, through intraperitoneal secretion, single chain variable fragments (scFvs) targeting OxLDL reduced foam cell formation in cholesterol-fed *Ldlr^-/-^*mice^27^.

In addition to the role of OxLDL in cardiovascular disease, there is emerging evidence that T regulatory cells (Tregs) may play key atheroprotective roles. Tregs are decreased in the peripheral blood of patients with atherosclerosis^28^, possibly due to Treg plasticity and dysfunctional immunosuppression^29^, suggesting that maintenance of local tolerance is essential for disease prevention. Tregs have been shown to inhibit foam cell formation *in vitro*^30^; additionally, in animal models, adoptive transfer of Tregs significantly increases IL-10 production while reducing both lesion size and IFN-γ production^31^.

Recent advances in adoptive cellular immunotherapy in cancer have ushered in new engineering platforms for personalized treatment. In particular, remarkable success has been achieved for patients with refractory and relapsed B cell leukemias and lymphomas, as well as myeloma^32–35^, through genetic modification of patient peripheral blood T cells with synthetic T cell receptors, known as chimeric antigen receptors (CARs), which utilize antibody-specificity to re-direct T cells to specific targets. This retooling of T cell specificity allows for immunoreactivity against self-antigens in addition to the response conferred by the endogenous T cell receptor (TCR). For B cell malignancies, patient T cells are engineered to ablate all cells expressing the pan-B cell antigen CD19, leading to prompt and often durable eradication of B cells in blood, bone marrow, and tissues^35^. The applicability of CAR T cell therapies has broadened beyond cancer to target HIV-1 infection^36,37^, opportunistic fungal infections^38^, and even refractory autoimmune diseases by eradicating pathogenic antibody-producing B cells^39^. CAR Treg engineering has also been developed in preclinical studies to treat various autoimmune diseases such as colitis and multiple sclerosis^40,41^, and phase II clinical trials using CAR Tregs are currently underway to treat solid organ transplant rejection^42^.

In this study, we hypothesized that combining the antigen specificity of CARs with the cardioprotective benefits of Treg cells would reduce the development of atherosclerosis. **Figure 1** depicts the potential therapeutic mechanism of this proposed therapy. In this report, we developed human and murine CAR Treg cells directed against OxLDL that demonstrated immunosuppressive capacity in vitro and ameliorated atherosclerosis *in vivo*.

**Figure 1.**
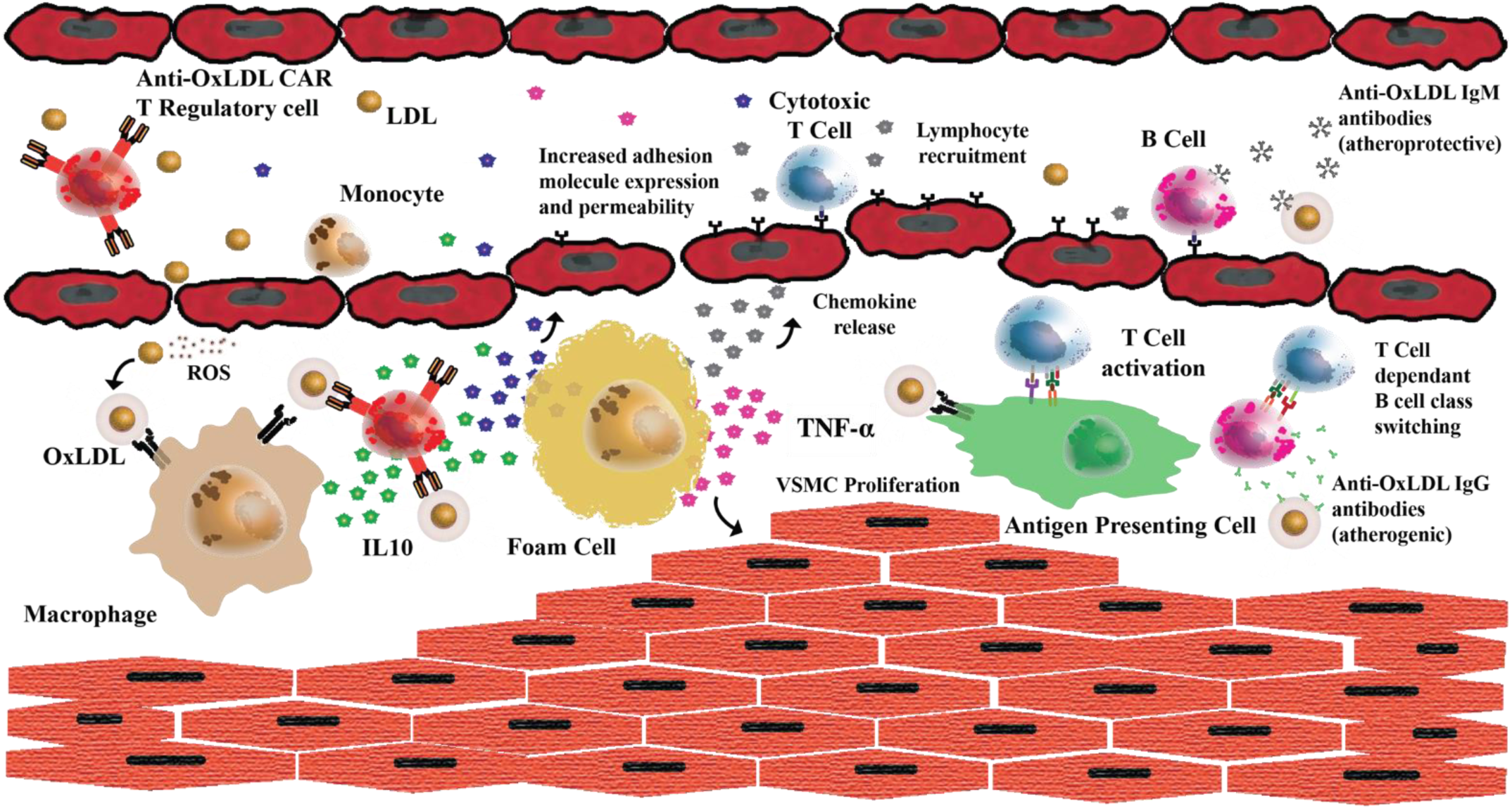
Pathogenesis of Atherosclerosis and Proposed anti-OxLDL CAR T Regulatory Therapy. LDL deposition in the endovascular endothelium results in oxidation by reactive oxygen species (ROS). OxLDL is internalized by scavenger receptors on macrophages, which turn into foam cells. Foam cells release pro-inflammatory cytokines (TNF-α) that increase adhesion molecule expression on the vascular endothelium and cause vascular smooth muscle cell (VSMC) proliferation. Chemokines are also released that recruit T cells and B cells. T cells become activated by foam cells or other antigen-presenting cells and then interact with B cells to produce pro-atherogenic anti-OxLDL immunoglobulins and increase inflammation. In time, a fibrous cap develops on the arterial wall, leading to plaque deposition and rupture. The addition of an anti-OxLDL CAR T regulatory cell therapy may reduce the inflammatory response initiated by OxLDL during the pathogenesis of atherosclerosis.

## Materials and Methods

### Vector Design

The cDNA sequence of the variable heavy and light chains of the IEI-E3 and 2D03 mAb were separated by a 3xG4S linker and custom synthesized by GeneArt (Life Technologies)^43^. The scFv was cloned into a CAR-encoding lentivirus backbone containing the CD8a leader sequence, a portion of the CD8a extracellular domain, CD28 transmembrane domain, and CD28 and CD3ζ endodomains. The human FOXP3 gene was inserted into the construct and separated from the CAR gene by a P2A peptide (**Figure 3A**). The final FOXP3-P2A-2D03-CD28-CD3z construct was inserted into the pTRPE vector: a custom synthesized third-generation lentiviral production vector using the EF1a promoter based on the similar pELNS vector previously described^44^. The murine FOXP3 gene, murine CD28, and murine CD3z replaced the human FOXP3, human CD28, and human CD3z genes for retroviral production. The entire murinized insert was cloned into the MSGV vector.

### Human T cell Transduction and Expansion

Lentiviral supernatant was generated from 293T cells transfected with pTRPE-IEI-E3-CD28-CD3z, pTRPE-2D03-CD28-CD3z, pTRPE-FOXP3-P2A-1E3-CD28-CD3z (non-specific CAR Tregs that targets an abhorrent glycosylation epitope of tumor-associated Tn antigen^45^), or pTRPE-FOXP3-P2A-2D03-CD28-CD3z plasmids, along with plasmids encoding gag/pol, Rev, and VSVg, as previously described^46^. Viral supernatants were collected and concentrated by ultracentrifugation at 24 hrs and 48 hrs post-transfection. Healthy donor CD4 T cells were positively selected from leukapheresis products using anti-CD4 (Miltenyi), activated in vitro with anti-CD3/CD28 magnetic beads (Life Technologies), and expanded in RPMI-1640 supplemented with 10% fetal calf serum (FCS), HEPES, penicillin and streptomycin (R10) and 200 IU/mL IL-2 for 10–15 days. T cells were transduced with lentiviral supernatant 16 hours after bead activation.

### Jurkat Stimulation Assay

NFAT-GFP reporter Jurkat cells were transduced with lentivirus encoding 2D03-CD28z and IEI-E3-CD28z CARs. Transduced Jurkat cells were normalized for CAR expression and then incubated for 24 hrs alone or with PMA/Ionomycin or plate-bound purified ApoB100, purified MDA-ApoB100, LDL, or MDA-LDL (Academy Bio-Medical Co.). Flow cytometry was then performed on Jurkat cells to evaluate GFP expression.

### Stimulation Assays

Non-transduced or transduced human T cells were incubated for 24 hours alone or on plates coated with OKT3 (anti-CD3), ApoB100, and MDA-ApoB100. The supernatant was removed to analyze for cytokine secretion. Cells were stained for surface and intracellular markers.

### Flow Cytometry

Anti-human antibodies were purchased from BioLegend, eBioscience, or Becton Dickinson. For cell surface staining, cells were washed with phosphate-buffered saline (PBS) containing 2% FCS and stained with Live/Dead Fixable violet (Thermo Fisher Scientific), and then stained for CAR, CD25, CD127, CD69, GARP depending on the experiment. For intracellular staining, cells were stained for viability and cell surface markers, fixed, permeabilized, and intracellularly stained for CTLA-4, Helios, and FoxP3. In all analyses, singlets were gated using FSC-H versus FSC-A and SSC-H versus SSC-A, followed by gating based on forward versus side scatter characteristics. Surface expression of 2D03-CAR was detected by staining with a biotin-conjugated goat anti-mouse F(ab)2 antibody (Jackson ImmunoResearch) and PE or APC-conjugated streptavidin. Flow cytometry was performed on a 4-laser LSRII or LSRFortessa (Becton Dickinson).

### Cytokine and Lipid Analysis

Supernatants were analyzed for IFN-γ using the Human IFN-γ DuoSet ELISA (R&D Systems). Human Luminex assay was performed according to the manufacturer’s instructions (Invitrogen and Millipore, respectively). Murine cytokines from in vivo experiments were measured via V-Plex according to the manufacturer’s instructions (MSD). Lipid measurements from in vivo experiments were measured indirectly through total cholesterol and HDL levels on a VET AXCEL Chemical Analyzer (Alfa Wassermann).

### Immunosuppression Assays

For in vitro suppression assays, 1×10^5^ human naïve CD4 and CD8 T cells were stained with CSFE, stimulated with anti-CD3 and anti-CD28 beads, and co-cultured with non-transduced CD4 T cells, 2D03-CAR Teffs or 2D03-CAR Tregs on MDA-ApoB100 coated 96-well plates. After 5 days, the samples were analyzed by flow cytometry, analyzing CSFE dilutions. Murine proliferation assays were performed using CSFE-stained murine 2D03-CAR Teffs stimulated with MDA-ApoB100 and analyzed after 5 days. Suppression assay was performed by adding murine 2D03-CAR Tregs at the indicated ratios.

### Human Macrophage Differentiation

Healthy donor monocytes were negatively selected from leukapheresis products using a RosetteSep kit (STEMCELL Technologies). Monocytes were differentiated to M0 with 10 ng/ml M-CSF for 1 week before use in subsequent experiments.

### Foam Cell Formation and Co-Culture Experiments

M0 macrophages were seeded at 2×10^5^ cells in 24 well plates. After 24 hours, 2×10^5^ stimulated or non-stimulated transduced CD4 T cells from the same donor are added to each well. Twenty-four hours after stimulation, 30 mg/mL of OxLDL was added to each well for 48 hours of incubation to allow foam cell formation. T cells were aspirated, macrophages were washed in PBS, fixed, stained with Oil Red O (ORO), and imaged. After imaging ORO was eluted with 100% isopropanol, the elution was transferred to a 96-well plate, and optical density at 500 nm was measured.

### Murine T cell Transduction and Expansion

Retroviral supernatant was generated from platE cells transfected with MSGV-muFOXP3-P2A-2D03-muCD28-muCD3z, MSGV-2D03-muCD28-muCD3z, MSGV-muFOXP3-P2A-1E3-muCD28-muCD3z, MSGV-CBG-T2A-GFP plasmids, plus gag/pol and collected at 48 hrs and 72 hrs. Retroviral supernatant was concentrated on retronectin-coated plates for 1 hour before spinfection of mouse CD4 T cells. Mouse CD4 T cells were isolated from bulk splenocytes by the EasySep Mouse CD4 T cell Isolation Kit (STEMCELL Technologies), activated with murine anti-CD3/CD28 beads (Thermo Fischer Scientific), and cultured with RPMI-1640 supplemented with 10% fetal calf serum (FCS), sodium pyruvate, β-Mercaptoethanol, penicillin and streptomycin (R10), and 200 IU/mL IL-2 for 5 days. T cells were transduced with retroviral supernatant 16 hours after bead stimulation.

### Murine Macrophage Differentiation

Total bone marrow cells were isolated from both the femora and tibiae of a C57BL/6 mouse as previously described^47^. Cells were suspended into RPMI1640 media supplemented with 10% heat-inactivated FCS and 100 ng/mL of murine M-CSF (complete media) at a density of 3 × 10^5^ cells/ml, seeded onto 12-well plates, and cultured for 5–7 days. The adherent cells were hereafter referred to as murine macrophages.

### Macrophage OxLDL Uptake Assay and Co-Culture Assay

In the OxLDL uptake assay, murine macrophages were seeded at 3 × 10^5^ cells/ml density in a 96-well plate with 50 ng/mL of LPS. After 24 hours, murine CD4 T cells, control Tregs, and 2D03-CAR Tregs were added at a density of 1 x 10^6^ cells/mL to each well, along with 50 ugl/mL of Dil-OxLDL. Dil-OxLDL uptake was measured via fluorescence over 60 hours on an xCELLigence Esight instrument (Agilent). In the co-culture experiment, murine macrophages were seeded at 3 × 10^5^ cells/mL density in 12-well MDA-ApoB100 coated plates. After 24 hours, the macrophages continued incubating alone, with control CAR Tregs or 2D03-CAR Tregs. After 24 hours, T cells were aspirated, macrophages were washed and detached with Accutase (Biolegend), stained for cell surface markers CD206, CD86, and MHCII, subsequently fixed with 4% PFA, and stained for intracellular marker Arginase.

### In Vivo Models

The experiments from this study were approved by the Institutional Animal Care and Use Committee at the University of Pennsylvania and were performed in accordance with Federal and Institutional Animal Care and Use Committee requirements. The T cell persistence experiment was conducted with male *C57BL/6J* mice. They were injected intraperitoneally (IP) with transduced T cells 24 hours after 150 mg/kg of cyclophosphamide or saline pre-conditioning (N = 8). Ten minutes before imaging, mice were injected with 3mg luciferase in 100 ul. Flux data was quantified using Living Image 4.5. Images were performed weekly until there was a lack of signal.

The Rader Lab previously developed *Ldlr*^-/-^/*Apobec1*^-/-^/human *ApoB* transgenic (LA-DKO/hApoB-Tg) mice used for in vivo studies^48^. At approximately 8-12 weeks of age, male and female mice (N=4) were conditioned with 150 mg/kg of cyclophosphamide via intraperitoneal injection. Approximately 24 hours after conditioning, the mice received intraperitoneal infusions of 1 x 10^7^ murine T cells (non-transduced CD4 T cells, control Tregs, or 2D03-CAR Tregs) and were started on a high-fat diet (Fisher Scientific). Transduced murine T cells were 30-40% CAR positive. Approximately 5 weeks after the first T cell injection, the mice received an additional round of cyclophosphamide conditioning and T cells. Five weeks after the final treatment (week 12), blood and serum were collected for lipid and V-Plex assay. After euthanasia, the mice were then perfused with PBS, and the heart and thoracic/abdominal aortas were removed and fixed for *en face* staining and histology.

### *En Face* Staining

Thoracic and abdominal aortas were dissected and fixed in 4% paraformaldehyde. After fixation, the aortas were opened, flattened, pinned, washed three times with PBS, once with 70% EtOH, and stained for 10 minutes with Sudan IV. Subsequently, stained aortas were washed once with 70% EtOH and imaged using a Nikon SMZ 1500 stereoscope with a Nikon Ri1 digital camera. The extent of plaques in the root or aorta or sclerosis in the root was quantified in ImageJ. The total Sudan IV staining portion was divided by the total area of the pinned descending aorta to obtain a proportion of the aorta involved by the lesion.

### Aortic Root Histopathology

Mouse hearts were fixed, dehydrated, embedded in paraffin, and serially sectioned. H&E and Masson’s Trichome staining was performed on paraffin-embedded tissues. Aortic root cross-sectional lesion areas were quantified using two representative cross-sections taken between the first appearance of the first leaflet of the aortic valve and the last booklet. The mean lesion size at each 100 -μm section in each animal was determined by computer-assisted morphometry (Image-Pro Plus 6.3, Media Cybernetics).

### Statistics

All in vitro experiments were performed in triplicate and independently at least twice. The results are shown as the mean ± SEM. One- or two-way paired ANOVA test was used to compare differences between groups in the various experiments. Differences of a *p*-value ≤ 0.05 were considered statistically significant.

## Results

### Design and construction of an anti-OxLDL CAR

Two CARs were developed from previously characterized anti-OxLDL antibodies, 2D03 and IEI-E3, which reduced atherosclerotic plaque formation when infused into *ApoE*^-/-^ mice by up to 50% and 36%, respectively^49^. 2D03 and IEI-E3 mAbs recognize malondialdehyde (MDA)-modified ApoB100, an epitope present on OxLDL *in vivo*^50^ and is commonly found in human atherosclerotic plaques^51^. ScFvs were generated by fusing the variable heavy and light chains of the 2D03 and IEI-E3 mAbs with the 15-mer (G4S)3 linker, and the scFvs were incorporated into CAR backbones containing CD8α hinge and transmembrane domain, CD28 intracellular signaling domain (ICD) and CD3ζ activation domain (**Figure 2A**). To efficiently assess which CAR would produce the most robust response to OxLDL and MDA-ApoB100, NFAT-GFP reporter Jurkat cells were transduced with lentiviral constructs encoding the 2D03 or IEI-E3 CARs. The transduced Jurkat cells were stimulated with native ApoB100, MDA-ApoB100, native LDL, and OxLDL. Both CARs exhibited NFAT signaling in response to MDA-ApoB100; however, the 2D03-28z CAR produced a significantly more robust NFAT signaling response to OxLDL (**Figure 2B**). Thus, the 2D03 CAR was chosen as the lead candidate molecule for future studies and referred to as 2D03-CAR.

**Figure 2.**
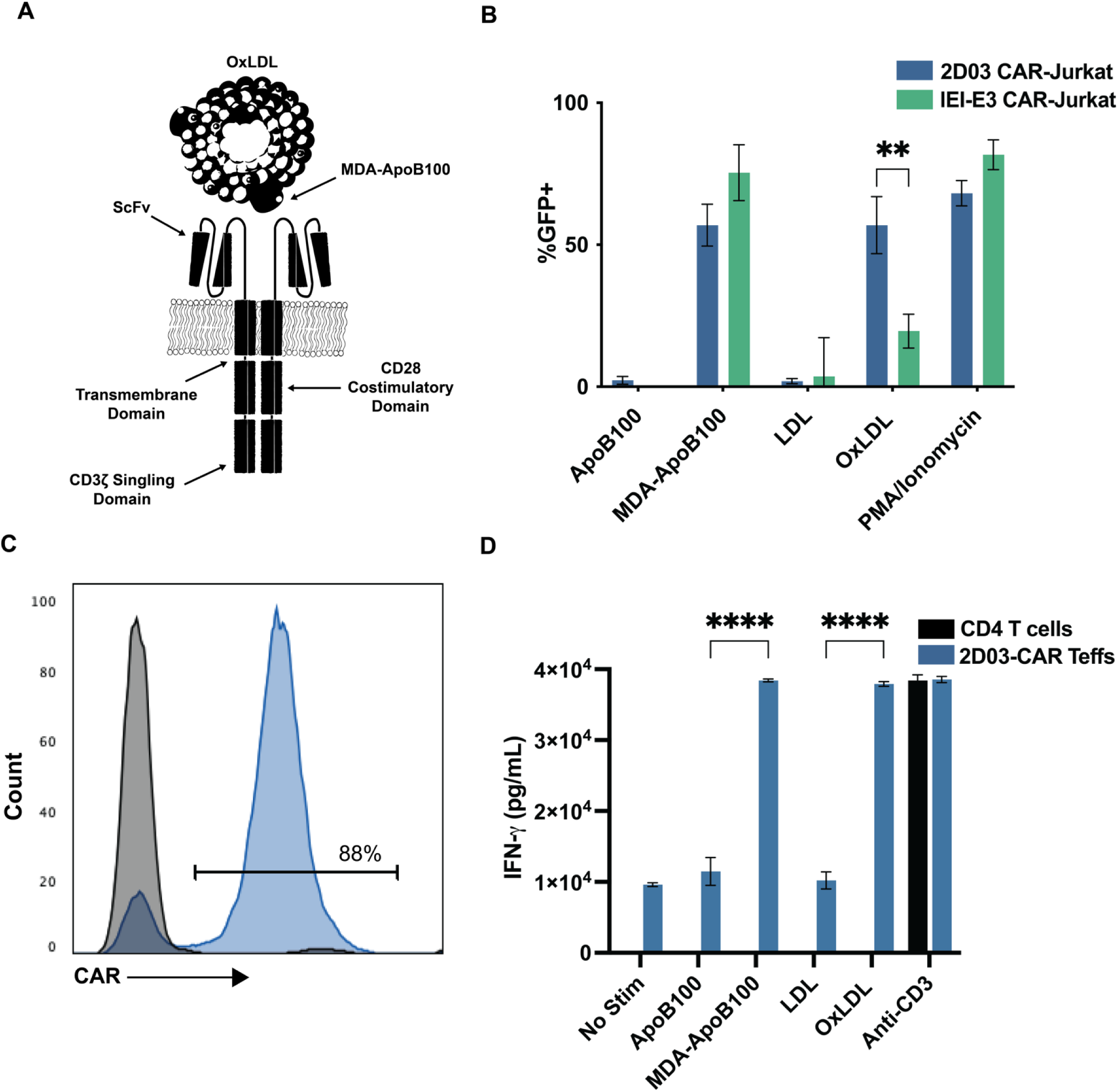
Anti-OxLDL CAR Design, Expression, and Specificity. **A)** Chimeric antigen receptor consists of a scFv, CD8α hinge and transmembrane domain, CD28 intracellular signaling domain, and CD3ζ endodomain. **B)** Stimulation assay based on percent change in GFP expression in stimulated NFAT-GFP reporter Jurkat cells. NFAT-GFP Jurkat cells transduced with lentivirus encoding 2D03 CAR or IEI-E3 CAR were cultured on ApoB100, MDA-ApoB100, LDL, or MDA-LDL coated plates for eight hours. PMA/Ionomycin was used as a positive control for cell stimulation. Statistical significance was calculated by comparing OxLDL-stimulated 2D03-CAR Jurkats with IEI-E3-CAR Jurkats. **C)** Flow cytometry histogram analysis of CAR^+^ T cells (blue) and non-transduced (NTD) T cells (black) stained with biotinylated Protein-L + streptavidin-PE. **D)** Indirect ELISA assays quantifying the IFN-γ production in supernatant from 2D03-CAR Teff cells and CD4 T cells cultured on OKT3 (anti-CD3), ApoB100, MDA-ApoB100, LDL, or MDA-LDL coated plates for 24 hrs. Protein/lipoprotein was plated at 10 μg/mL. Statistical significance was calculated comparing cytokine production of 2D03-CAR Teffs cells when exposed to ApoB100 vs. MDA-ApoB100 and LDL vs. MDA-LDL. ✱✱= p ≤ 0.01, ✱✱✱✱ = p ≤ 0.0001.

Next, we stimulated normal donor CD4 T cells with anti-CD3 and anti-CD28 mAb-coated magnetic beads and transduced them with a lentiviral vector encoding the 2D03-CAR. After transduction and expansion, 70-90% of T cells express the CAR (**Figure 2C**). To evaluate the specificity of the 2D03-CAR on T cells, non-transduced CD4 T cells and 2D03-CAR T cells were incubated on plates coated with the anti-CD3 mAb (OKT3), ApoB100, MDA-ApoB100, native LDL, and OxLDL for 24 hours. The culture supernatants were assessed for IFN-γ by ELISA (**Figure 2D**). 2D03-CAR T effector cells (2D03-CAR Teffs) secreted similar IFN-γ quantities in response to the OxLDL, MDA-ApoB100, and anti-CD3, but not native LDL or ApoB100. The production of IFN-γ by helper CD4 T cells demonstrates the capability of oxidized lipoproteins to stimulate CAR T cells, and the lack of 2D03-CAR T cell stimulation by native LDL also reflects the conservation of specificity of the 2D03-CAR for oxidation-specific epitopes.

### Constitutive FoxP3 expression in CD4 T cells induces T regulatory phenotype and function

To develop Tregs, human peripheral blood CD4 T cells were transduced with a bicistronic lentiviral vector that co-expresses FoxP3 and CAR separated by a P2A ribosomal skip element (CAR-Treg vector) (**Figure 3A**). This method has been previously shown to develop stable Tregs^52^. T cells transduced with the CAR-Treg vector were evaluated for FoxP3 expression, which was significantly higher in the T cells transduced with the CAR-Treg vector than cells transduced with the CAR-Teff vector, confirming dual expression from the bicistronic vector (**Figure 3B**). Common Treg surface markers, such as CD127, CD25, and CTLA-4, were assessed by staining the FoxP3^+^CAR^+^ CD4 T cells (2D03-CAR Tregs) and compared to unmodified CD4 T cells, 2D03-CAR Teffs, and endogenous Treg cells obtained from the same donor. 2D03-CAR Tregs were phenotypically CD25^hi^ and CD127^low^ and exhibited similar levels of CTLA-4 compared to endogenous Tregs (**Figure 3C**). Upon stimulation of 2D03-CAR Teffs and 2D03-CAR Tregs with anti-CD3, 2D03-CAR Tregs cells did not produce a significant increase in IFN-γ production in contrast to the 2D03-CAR Teffs (**Figure 3D**), demonstrating the effect of FoxP3 expression on inhibition on inflammatory cytokine production. 2D03-CAR Teffs and 2D03-CAR Tregs were co-cultured with naïve T cells in the presence of anti-CD3 and anti-CD28 beads, and the proliferation of naïve T cells was measured by CSFE dilution to determine the immunosuppressive capacity of 2D03-CAR Tregs on activated T cells. (**Figure 3E**). While non-transduced CD4 T cells and 2D03-CAR Teffs had minimal effects on the proliferation of naïve T cells, 2D03 CAR-Tregs significantly reduced the proliferation of naïve T cells, demonstrating gained suppressive function through TCR stimulation; thus, ectopic expression of FoxP3 in CAR+ CD4 T cells induced a Treg phenotype and function.

**Figure 3.**
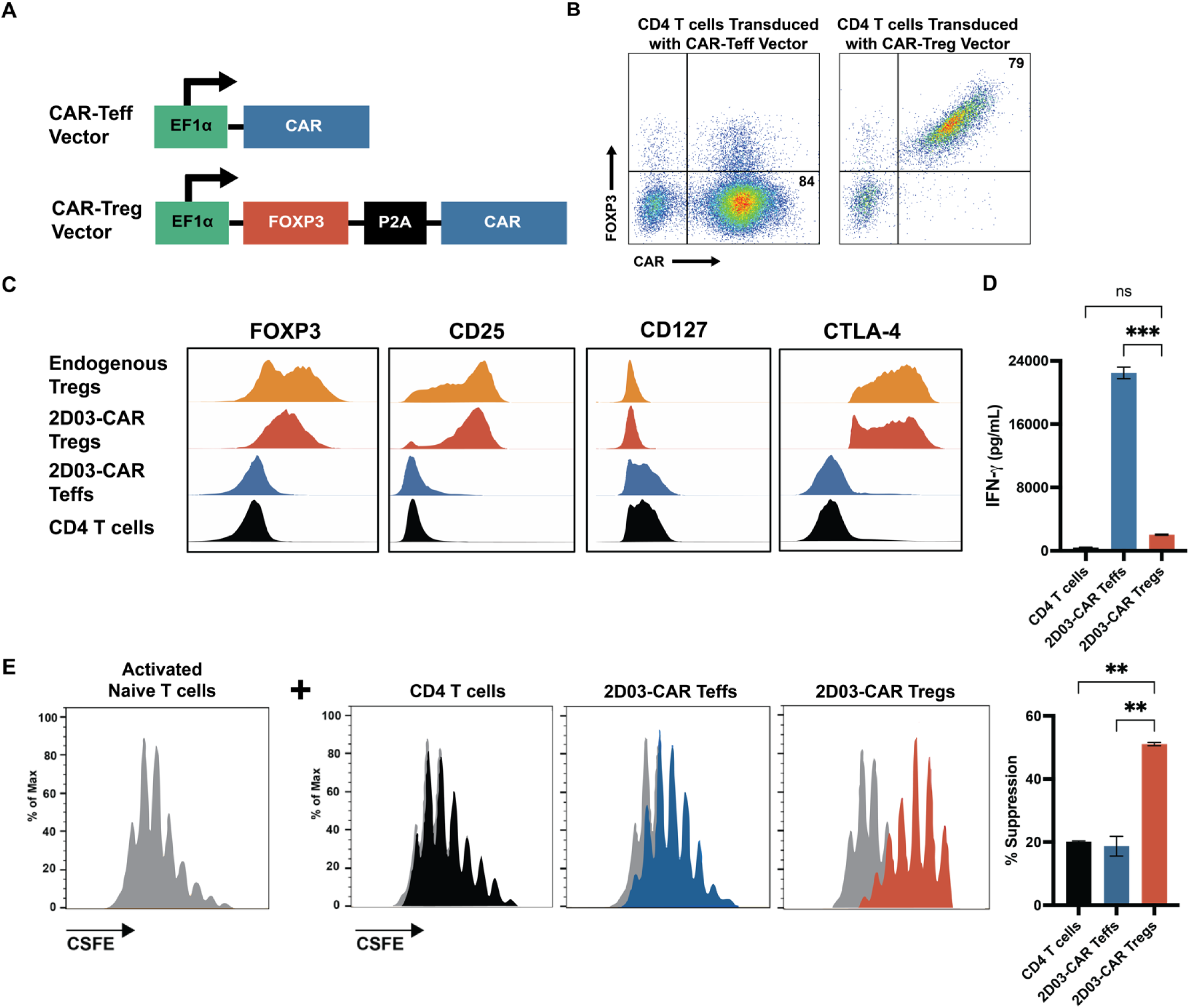
T regulatory Induction, Phenotype, and Behavior. **A)** The CAR-Treg vector contains an EF1α promotor followed by the human *FoxP3* transgene; the CAR was inserted after a P2A peptide sequence. The entire expression cassette is translated into a FoxP3-P2A-CAR fusion protein separated at the P2A element to produce the two separate proteins, FoxP3 and CAR. **B)** Intracellular staining assay measuring expression of FoxP3 and CAR in CD4 T cells transduced with the CAR-Teff vector and the CAR-Treg vector. **C)** Comparison of FOXP3, CTLA-4, CD25, and CD127 staining of CD4 T cells, 2D03-CAR Teffs, 2D03-CAR Tregs, and endogenous Tregs from the same donor. **D)** Indirect ELISA assays quantifying the IFN-γ production in supernatant from CD4 T cells, 2D03-CAR Teffs, and 2D03-CAR Tregs cultured on MDA-ApoB100 coated plates for 24 hours. Statistical significance was calculated by comparing cytokine production of 2D03-CAR Tregs to 2D03-CAR Teffs and CD4 T cells. **E)** Immunosuppression assay of CSFE-labeled naïve human T-cells activated with anti-CD3/anti-CD28 paramagnetic beads and cultured alone or in the presence of CD4 T cells, 2D03-CAR Teffs or 2D03-CAR Tregs. Percent suppression was calculated using index divisions of stimulated naïve T cells in treatment groups compared to those of stimulated naïve T cells alone. Statistical significance was calculated by comparing the percent suppression of naïve T cells when incubated with 2D03-CAR Tregs vs CD4 T cells or 2D03-CAR Teffs. ✱✱= p ≤ 0.01, ✱✱✱ = p ≤ 0.001.

### 2D03-CAR Tregs retain immunosuppressive phenotype in vitro when stimulated through the CAR or TCR and reduce foam cell formation

To investigate whether 2D03-CAR Tregs retain their phenotype after stimulation through CAR, non-transduced CD4 T cells and 2D03-CAR Tregs were incubated alone or on plates coated with MDA-ApoB100 or anti-CD3 and stained for various Treg and T cell activation markers 24 hours after culturing (**Figure 4A**). 2D03-CAR Tregs retained a CD25^hi^CD127^low^ phenotype after stimulation. Similarly, Treg activation markers, such as GARP, CTLA-4, and Helios, increased in CAR Tregs upon stimulation through the CAR, along with the T cell activation marker, CD69. Cytokine production of 2D03-CAR Tregs was measured from culture supernatant after stimulation by the CAR antigens (**Figure 4B**). 2D03-CAR Tregs produced anti-inflammatory cytokines IL-13, IL-10 and IL-4, along with increased production of IL-2R. Thus, stimulation of 2D03-CAR Tregs through the CAR retains the Treg phenotype and demonstrates the capacity for contact-dependent immunosuppression through increased inhibitory surface markers, such as CTLA-4, and contact-independent immunosuppression via anti-inflammatory cytokines secretion.

**Figure 4.**
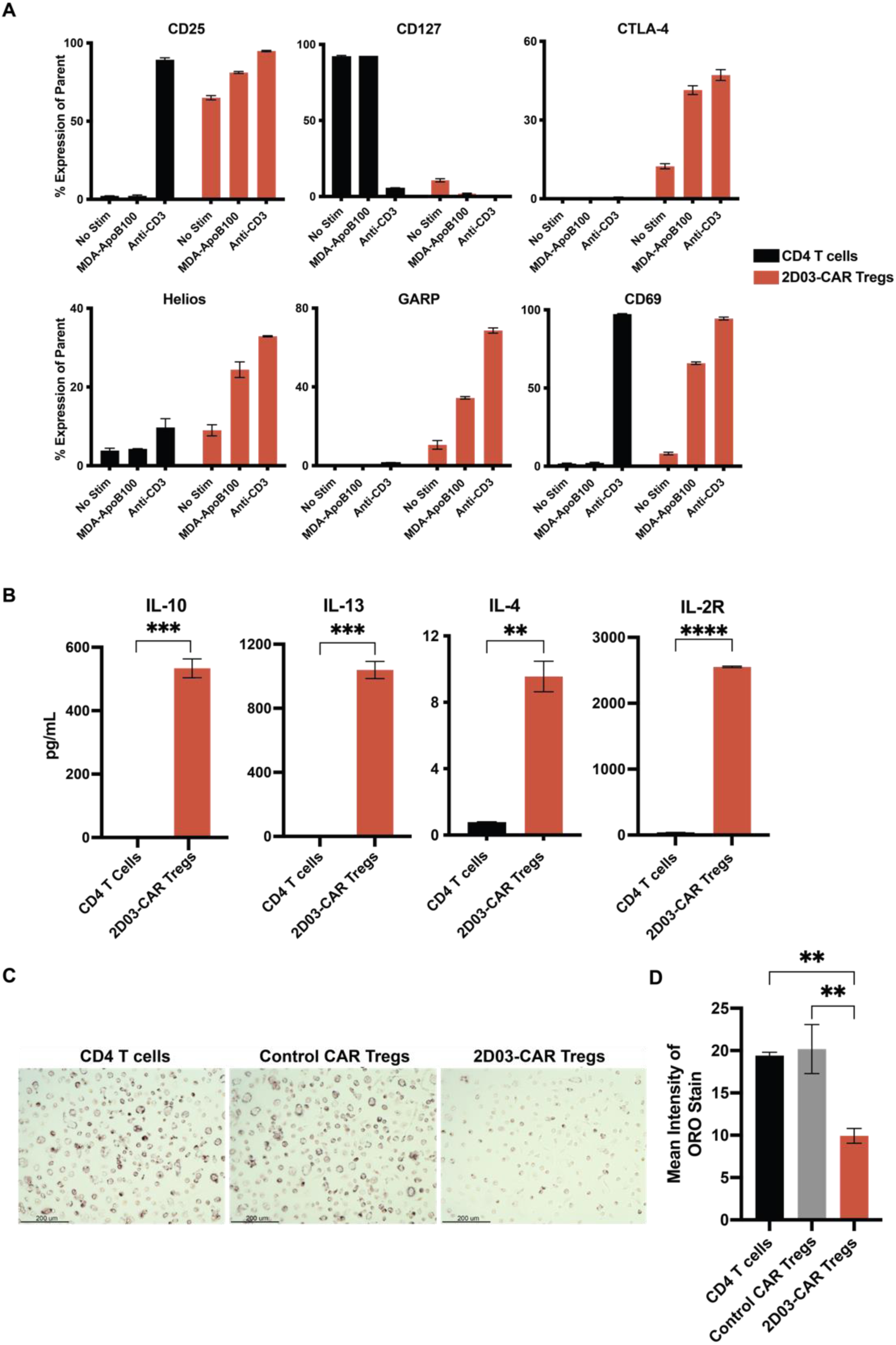
2D03-CAR Tregs Plasticity, Cytokine Production, and Macrophage Foam Cell Formation. **A)** Stimulation assay quantifying percent of cells expressing respective markers. CD4 T cells or 2D03-CAR Tregs were cultured with no stimulation, MDA-ApoB100, or anti-CD3 coated plates for 24-hrs, and then stained for both cell surface and intracellular markers. **B)** Indirect ELISAs quantifying IL-10, IL-13, IL-4, and IL-2R production in supernatant from the stimulation assay. Statistical significance was calculated by comparing the cytokine production of CD4 T cells to 2D03-CAR Tregs when exposed to MDA-ApoB100. **C)** Representative images of Oil Red O (ORO)-stained foam cells. 2×10^5^ macrophages were seeded on a 24-well plate coated with 10mg/ml of MDA-ApoB100 and non-transduced CD4 T cells, control CAR Tregs, or 2D03-CAR Tregs of the same normal donor. After 48 hours, 50mg/ml of MDA-LDL was added to the culture to induce foam cell formation. Forty-eight hours later, the macrophages were fixed and stained with ORO. **D)** Relative intensity of ORO staining in Fig. 2C. Statistical significance was calculated by comparing macrophages cultured with 2D03-CAR Tregs to control groups. ✱✱= p ≤ 0.01, ✱✱✱ = p ≤ 0.001, ✱✱✱✱ = p ≤ 0.0001.

We next asked whether 2D03-CAR Tregs could reduce foam cell formation, given its clinical importance in the generation of atherosclerosis. Macrophages were incubated on plates coated with MDA-ApoB100 alone, with CD4 T cells, control CAR Tregs (expressing a non-specific CAR), or 2D03-CAR Tregs. After 48 hours, OxLDL was added to the culture to induce foam cell formation. Oil Red O (ORO) staining of the macrophages 48 hours after OxLDL addition demonstrated a ∼50% reduction in lipid uptake in macrophages incubated with the 2D03-CAR Tregs compared to all other treatment groups (**Figure 4C** and **4D**). All other groups had comparable amounts of lipid uptake, demonstrating that OxLDL-specific Tregs, but not non-specific Tregs, are sufficient to reduce foam cell formation.

### Murine 2D03-CAR Treg development and characterization

Next, we established and characterized murine versions of the 2D03-CAR Tregs, in which human FOXP3 and the human CD28 signaling domain were replaced with homologous murine sequences. This cassette was inserted into a gamma retroviral vector, which exhibits enhanced transduction efficiency in murine T cells compared to lentiviral vectors^53^. Given the prior efficacy of the 2D03 antibody in murine models^41,54^, no other alterations were made to the CAR. The murine 2D03-CAR Tregs were CD25^hi^CD127^low^, suppressed murine 2D03-CAR Teffs in response to MDA-apoB100 stimulation, and persisted for up to 5 weeks in *C57BL/6J* mice when injected intraperitoneally 24 hours after cyclophosphamide pre-conditioning (**Supplementary Fig. 1-2**).

### Murine OxCAR Tregs reduce OxLDL uptake and increase arginase expression

To assess whether the murine 2D03-CAR Tregs can similarly reduce OxLDL uptake, we incubated Lipopolysaccharide (LPS)-stimulated murine macrophages with fluorescently labeled OxLDL (Dil-OxLDL) and either murine non-transduced CD4 T cells, control CAR Tregs, or 2D03-CAR Tregs and subsequently measured fluorescent uptake over time (**Figures 5A** and **5B**). Uptake of Dil-OxLDL increased over time in each group; however, the presence of 2D03-CAR Tregs significantly reduced total uptake compared to other groups at 60 hours.

**Figure 5:**
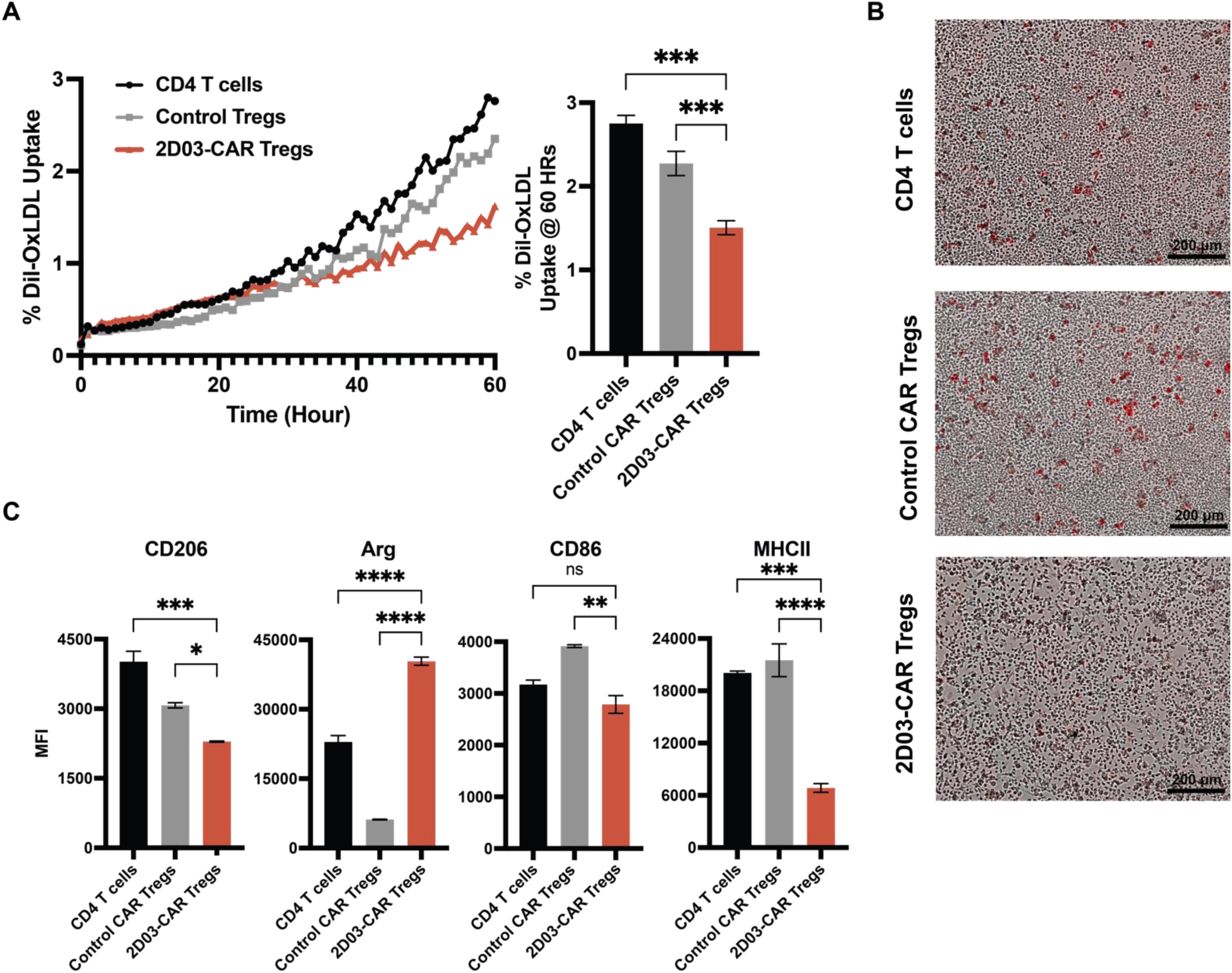
Murine Macrophage Uptake of Dil-OxLDL and Phenotype. **A)** Murine macrophages were co-cultured with CD4 T cells, control CAR Tregs, and 2D03-CAR Tregs in the presence of LPS and Dil-OxLDL for 60 hours. Dil-OxLDL uptake was measured by fluorescence signal. Statistical significance was calculated by comparing Dil-OxLDL at 60 hrs in macrophages co-cultured with 2D03-CAR Tregs to control groups. **B)** Representative images of Dil-OxLDL uptake between each treatment group during the experiment. **C)** Cell surface and intracellular staining of murine macrophages incubated with CD4 T cells, control CAR Tregs, or 2D03-CAR Tregs for 24 hours. Statistical significance was calculated by comparing the mean fluorescent intensity (MFI) of the surface or intracellular marker on macrophages co-cultured with 2D03-CAR Tregs to control groups ✱= p ≤ 0.05, ✱✱= p ≤ 0.01, ✱✱✱ = p ≤ 0.001, ✱✱✱✱ = p ≤ 0.0001.

We hypothesized that the reduction in uptake may be due to macrophages skewing towards an anti-inflammatory phenotype. To assess the influence of murine 2D03-CAR Tregs on macrophage phenotype, murine macrophages were incubated alone, with control CAR Tregs, or stimulated 2D03-CAR Tregs and stained for M1 and M2 markers (**Figure 5C**). Although there was a reduction in the mean fluorescent intensity (MFI) of CD206 on macrophages incubated with 2D03-CAR Tregs, we noted significantly decreased MHCII expression and markedly increased arginase expression, consistent with an anti-inflammatory phenotype. These data suggest that activated 2D03-CAR Tregs limit foam cell formation and skew macrophages towards an anti-inflammatory phenotype.

### Murine 2D03-CAR Tregs reduces atherosclerosis in vivo without systemic change in inflammatory markers

*Ldlr*^-/-^;*Apobec1*^-/-^;human *ApoB* transgenic (LA-DKO/hApoB-Tg) mice lack the *Apobec1* gene responsible for editing full-length ApoB100 mRNA in liver to generate the ApoB48 protein and thus express only full-length human apoB100 in liver (similar to humans) at high levels; these mice have 3-fold higher plasma levels of cholesterol than *Ldlr*^−/−^ mice and develop extensive atherosclerotic lesions with human physiologic levels of apoB100 expression^49^. We evaluated whether murine 2D03-CAR Tregs can reduce atherosclerotic plaque burden *in vivo* in LA-DKO/hApoB-Tg mice. We treated 8-12 week-old LA-DKO/hApoB-Tg mice (conditioned 24 hours prior with cyclophosphamide) with murine non-transduced CD4 T cells, control CAR Tregs, or 2D03-CAR Tregs. The mice were started on a high-fat diet one day after T cell treatment and received a second injection of T cells five weeks after the first T cell injection, given our prior data (Supp. Fig. 2) demonstrating the persistence of murine CAR T cells to be around 6 weeks, consistent with other studies^46^. After two and a half months, the mice were euthanized, and serum, hearts, and aortas were harvested for analysis. The descending aortas of each mouse were analyzed for atherosclerotic lesions via *en face* staining. The mice receiving the 2D03-CAR Tregs had a mean plaque burden of 1.38% ± .29%, which was significantly reduced compared to mice treated with CD4 T cells (7.13% ± .84%; p = .05) and control CAR Tregs (8.70% ± 1.86%; p = .02) (**Figure 6A** and **6B**). Compared to control CAR Tregs, the 2D03-CAR Tregs resulted in a mean reduction of plaque burden of nearly 80% (**Figure 6C**). Notably, serum lipid analysis showed no significant changes in LDL levels among treatment groups (**Figure 6D**), indicating that the reduced atherosclerosis was not due to systemic reductions in LDL, distinguishing the mechanism of action 2D03-CAR Tregs from lipid-lowering drugs.

**Figure 6:**
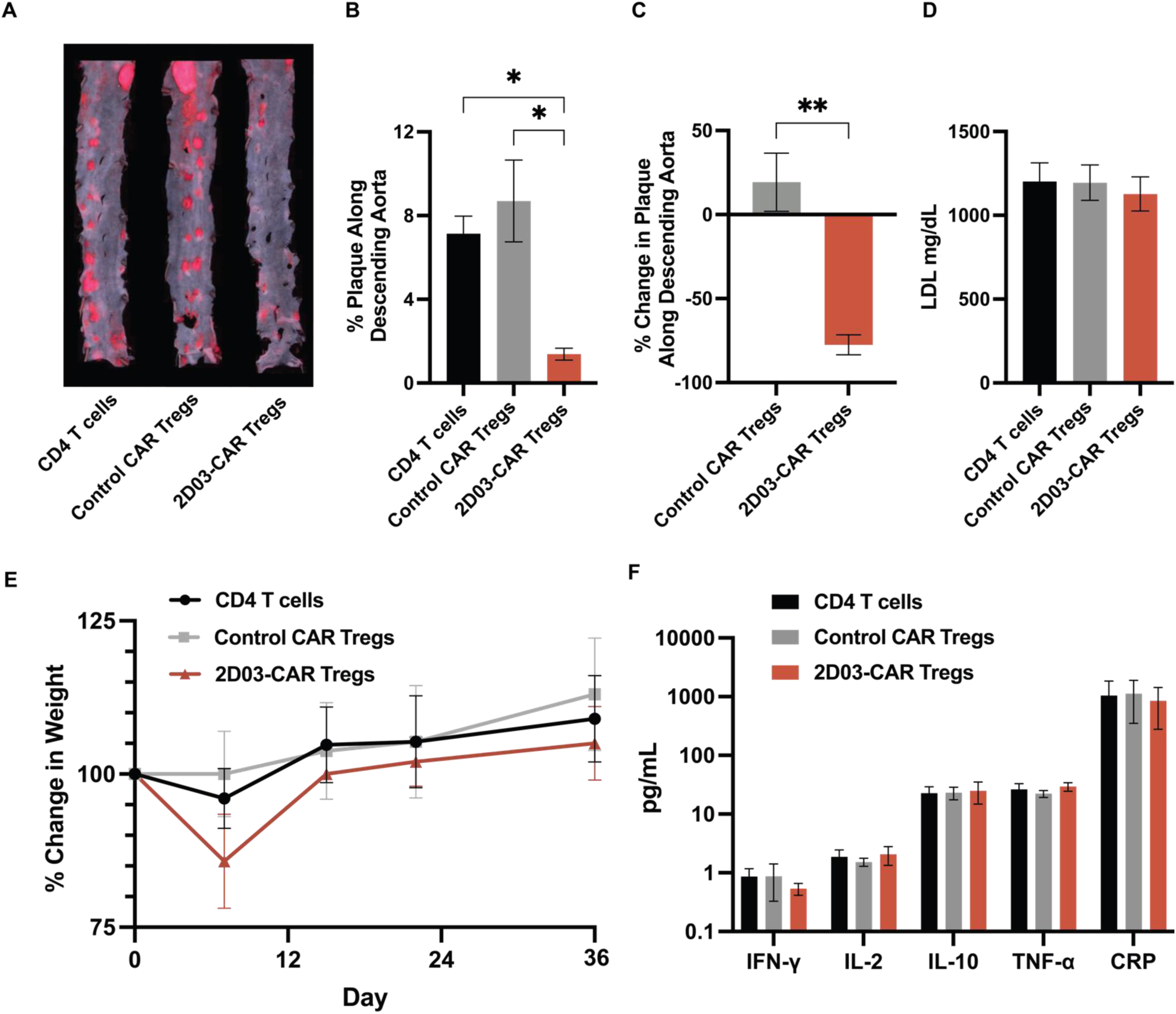
Safety and efficacy of 2D03-CAR Tregs in hyperlipidemic mice. **A)** Representative images of Sudan IV staining of descending aortas. **B)** Quantification of the percentage of plaque along the total descending aorta in each treatment group. Statistical significance was calculated by comparing the plaque burden percentage of mice treated with 2D03-CAR Tregs to control groups. **C)** Average percent change in plaque burden in mice treated with either control CAR Tregs or 2D03-CAR Tregs compared to baseline (CD4 T cell treated mice). Statistical significance was calculated by comparing the change in percentage of plaque burden of mice treated with 2D03-CAR Tregs compared to control CAR Tregs. D) Average serum LDL levels in respective treatment groups at the end of the experiment. No significant difference was observed among treatment groups. **E)** Percent change in body weight of mice during the first month of treatment with CD4 T cells, control CAR Tregs, and 2D03-CAR Tregs. No significant weight difference was observed among treatment groups. **F)** Serum cytokines of each treatment group at the end of the experiment. No significant differences were noted. ✱= p ≤ 0.05, ✱✱= p ≤ 0.01.

We also observed no significant toxicity in mice treated with 2D03-CAR Tregs. Although we observed a reduction in weight among each group 7 days after conditioning and treatment, the differences between treatment groups were not significant, and all mice re-gained weight at similar rates during the first month of treatment (**Figure 6E**). Finally, we did not observe significant differences in inflammatory or anti-inflammatory cytokines among treatment groups, suggesting a localized effect of the 2D03-CAR Tregs (**Figure 6F**).

Lastly, immunohistochemistry with H&E and Masson’s Trichome was performed on the aortic roots of each mouse. In accordance with the observation of decreased plaque in the descending aorta of the mice treated with the 2D03-CAR Tregs, there was a significant reduction in atherosclerotic plaque along the aortic root in mice treated with 2D03-CAR Tregs (**Figures 7A** and **7B**). Masson’s trichome staining demonstrated increased collagen content and aortic root in the mice treated with 2D03-CAR Tregs compared to control groups (**Figures 7C** and **7D**), suggesting an increase in plaque stability in 2D03-CAR Treg-treated mice.

**Figure 7:**
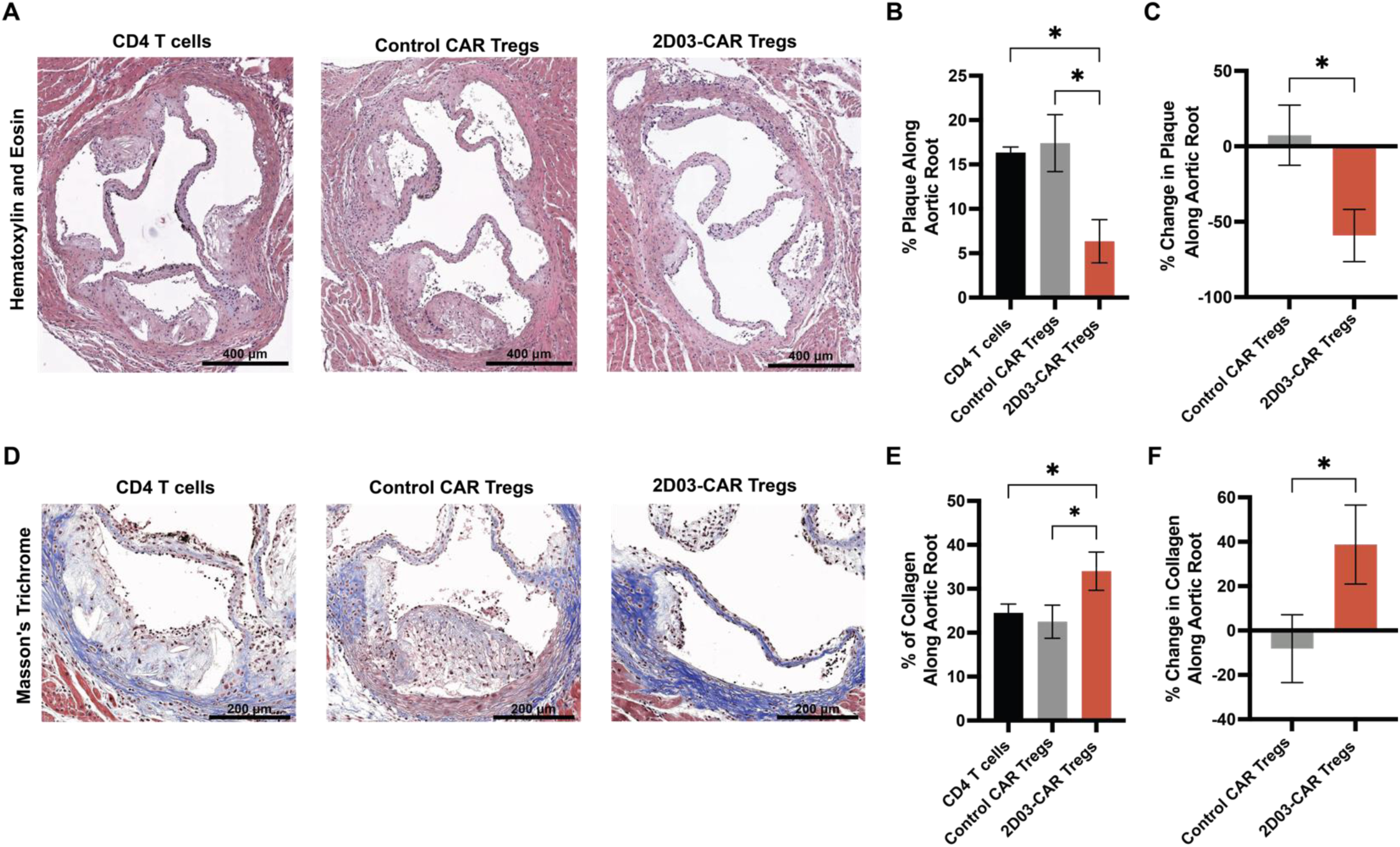
Immunohistochemical analysis of hyperlipidemic mice treated with 2D03-CAR Tregs. **A)** Representative images of H&E staining of aortic root. **B)** Quantification of plaque percentage in the aortic root. Statistical significance was calculated by comparing the percentage plaque burden along the aortic root of mice treated with 2D03-CAR Tregs to control groups. **C)** Average percent change in plaque burden along aortic root in mice treated with either control CAR Tregs or 2D03-CAR Tregs compared to baseline (CD4 T cell treated mice). Statistical significance was calculated by comparing the change in plaque burden percentage along the aortic root of mice treated with 2D03-CAR Tregs compared to control CAR Tregs. **D)** Representative images of Masson’s Trichrome staining of aortic root. **E)** Quantification of the collagen percentage along the aortic root. Statistical significance was calculated by comparing the percentage of collagen along the aortic root of mice treated with 2D03-CAR Tregs to control groups. **F)** Average percent change in collagen percentage along the aortic root in mice treated with either control CAR Tregs or 2D03-CAR Tregs compared to baseline (CD4 T cell treated mice). Statistical significance was calculated by comparing the change in collagen content percentage of mice treated with 2D03-CAR Tregs compared to control CAR Tregs. ✱= p ≤ 0.05.

## Discussion

Here, we report the first application of CAR T cell technology to target atherosclerotic cardiovascular disease. Murine 2D03-CAR Tregs directed towards a malondialdehyde-modified apolipoprotein B-100 epitope substantially reduced atherosclerosis in an atherosclerosis-prone mouse model. This demonstrates the feasibility of utilizing CARs to target epitopes not restricted to the cell surface and suggests that CAR Tregs may be a novel approach to target residual risk in atherosclerotic cardiovascular disease.

Dysfunctional immune tolerance and a skewed balance between pro-inflammatory T cells and functional regulatory T cells contribute to atherosclerosis^55^. Identifying pathogenic post-translational modifications of LDL has provided potential therapeutic targets to reduce atherosclerotic plaque size and associated inflammation.

Recent advances in cellular engineering now provide tools and platforms to synthetically re-target and re-equip the immune system with either cytotoxic or immunosuppressive properties and allow the generation of novel cellular drugs to treat pro-inflammatory diseases like atherosclerosis. In this report, we developed an anti-inflammatory cellular therapy targeting OxLDL to reduce atherosclerosis *in vivo*.

Clinically, Tregs are detected in lower frequencies in atherosclerotic plaques (1–5% of all T cells) than in other chronically inflamed tissues (∼25% of all T cells in eczema or psoriasis), which suggests that impairment of local tolerance is a cause for inflammation and plaque formation^56^. Immune intolerance to OxLDL contributes to atherosclerotic plaque development, and continual inflammation at these plaques from helper and cytotoxic T cells is one of the key drivers leading to plaque instability and rupture. Developing a method to induce immune tolerance to OxLDL at plaques through Treg induction, either via adoptive transfer and/or redirection, may provide a means to stabilize or cause regression of plaques. Not surprisingly, the potential suppressive effects of Tregs for inflammation have provoked an interest in clinical translation, particularly for autoimmune diseases. However, Tregs have only been adoptively transferred clinically to induce transplant tolerance^57^. The lack of clinical implementation of adoptive Treg therapies is likely due to the challenge of sorting and expanding natural Tregs since FoxP3, an intracellular transcription factor, is the only reliable marker. Furthermore, ensuring the stability of antigen-specific Tregs is another hurdle potentially preventing Tregs from broader clinical applications^58^. Prior studies have shown that this obstacle can be overcome with gene transfer of FoxP3 into CD4 T cells^52^; this methodology was applied to developing OxLDL-specific Tregs.

Human CD4 T cells were modified using lentivirus to co-express FoxP3 and CAR to guarantee that all CAR^+^ T cells were suppressive, preventing potential Treg plasticity^59^. The engineered Tregs expressed CAR and FoxP3 and resembled natural Tregs based on classic phenotypic markers. Regardless of stimulation through the natural TCR or the ectopically expressed CAR, 2D03-CAR Tregs maintained T regulatory phenotype, specificity, and suppressive capacity. In assays testing 2D03-CAR Treg function, the 2D03-CAR Tregs produced anti-inflammatory cytokines upon stimulation through CAR, significantly decreasing the proliferation of activated T cells. Most importantly, 2D03-CAR Tregs reduced foam cell formation when cultured with macrophages and OxLDL, previously shown to occur with polyclonal Tregs^60^.

We further developed and characterized murine 2D03-CAR Tregs. These Tregs demonstrated suppressive activity in vitro, decreasing macrophage uptake of OxLDL and increasing macrophage arginase-I expression. Arginase-1 expression in macrophages is associated with an anti-inflammatory (M2-like) phenotype^61^, supporting our hypothesis that the 2D03-CAR Tregs skew macrophages towards an anti-inflammatory phenotype. Furthermore, arginase-1 expression in macrophages has been associated with enhanced atherosclerotic plaque stabilization^62^. This may explain our observation of increased collagen content along the aortic roots in mice treated with 2D03-CAR Tregs.

Finally, we tested the efficacy of murine 2D03-CAR Tregs *in vivo*. Multiple considerations were needed to effectively test CAR T cell therapy in an atherosclerotic model. Standard *in vivo* models of atherosclerosis rely upon immunocompetent mice to produce the inflammatory milieu; however, most CAR T cell therapy models are performed in immunocompromised mice to enhance human CAR T cell persistence^63^. To enhance murine CAR T cell persistence in syngeneic models, lymphodepletion is completed via either chemotherapeutic conditioning or irradiation. We favored conditioning with cyclophosphamide for our preliminary experiment as the latter has been demonstrated to increase atherosclerotic burden in hyperlipidemic mice^64^. We additionally treated mice with multiple doses to ensure the survival of murine CAR T cells along the timeline of the experiment to achieve a therapeutic effect. Lastly, we used mice genetically engineered to express the full human ApoB100 transgene, increasing the translational utility of the 2D03-CAR Tregs. In the mice treated with 2D03-CAR Tregs, we observed a significant inhibition of atherosclerotic burden along the descending aorta and aortic root without substantial changes in systemic cytokines. These observations indicate that if the treatment had any effect, it may have been localized and not produced systemic inflammatory changes, or changes were transient and not present at the time of harvest. We additionally noted increased collagen content along the aortic root of mice treated with 2D03-CAR Tregs, which has previously been shown to be a surrogate of plaque stability^65^.

Our study has multiple limitations. We had a limited number of mice per treatment group and, therefore, acknowledge that applying parametric tests may violate the assumption of mean population normality. We did not assess atherosclerosis before treatment; thus, while atherosclerosis progression was reduced, we have not yet demonstrated regression of pre-existing disease. Finally, the clinical translation of murine 2D03-CAR Tregs is limited by differences between murine and human CAR T cell behavior, particularly in the duration of murine CAR T cell survival and longevity. Therefore, generating an alternative humanized model of atherosclerosis would be necessary to support the preclinical development and potential translation of human 2D03-CAR Tregs.

This is the first report of CAR T cell technology to reduce atherosclerosis, representing the first instance of cellular immunotherapy explicitly developed for atherosclerosis. Unlike current therapies that primarily target cholesterol levels or require invasive procedures, this approach provides a targeted, immune-mediated treatment that acts locally within atherosclerotic plaques. By engineering CAR Tregs to specifically recognize OxLDL, these cells can target atherosclerotic lesions and exert their immunosuppressive effects precisely at the disease site, reducing systemic side effects and directly addressing the underlying inflammatory pathways crucial to disease progression.

This type of treatment has significant therapeutic potential, particularly for patients who are refractory or have contraindications to standard management, such as patients with malignancy-associated coronary artery disease who face particular challenges due to increased risks of bleeding from therapies like aspirin or anti-coagulation and heightened rates of stent thrombosis after percutaneous interventions. Although other anti-inflammatory immunotherapies for cardiovascular disease have shown efficacy in reducing cardiac events, such as canakinumab in the CANTOS trial, many of these medications are associated with significant side effects or increased risk of higher incidence of fatal infections due to immunosuppression^66–68^. A targeted anti-inflammatory therapy like OxLDL-specific CAR Treg cells poses significantly less risk of these complications than invasive procedures and may be less systemically immunosuppressive. Taken together, the innovative aspects of this study—from CAR Treg engineering to targeted, local immune modulation—represent a significant advancement in cardiovascular disease treatment. The development of OxLDL-specific CAR Tregs provides a unique opportunity to shift the treatment paradigm from invasive and non-specific interventions to precise and effective cellular immunotherapy.

Considering that the manufacturing of CAR T cells is already functional globally and has shown clinical success in cancer^69^, the translation of 2D03-CAR Treg therapy could be easily implemented. Autologous CD4 T cells were isolated from the peripheral blood of normal donors and transduced with lentivirus to co-express FoxP3 and 2D03-CAR. 2D03-CAR Tregs may be a living drug that reverses and prevents atherosclerosis and heart disease when infused into the patient. Although other novel immunotherapies, such as anti-OxLDL immuno-cytokines, are currently being evaluated, there are many limits to these antibody therapies since they require multiple infusions, and there is an increased risk for hypersensitivity reactions^70^. 2D03-CAR Treg treatment, on the other hand, may provide a one-time injection with lifelong benefits through the development of OxLDL-specific memory Treg cells. The novel findings in this report warrant further investigation due to their significant clinical potential.

## Supplementary Figures

**Supplementary Figure 1:**
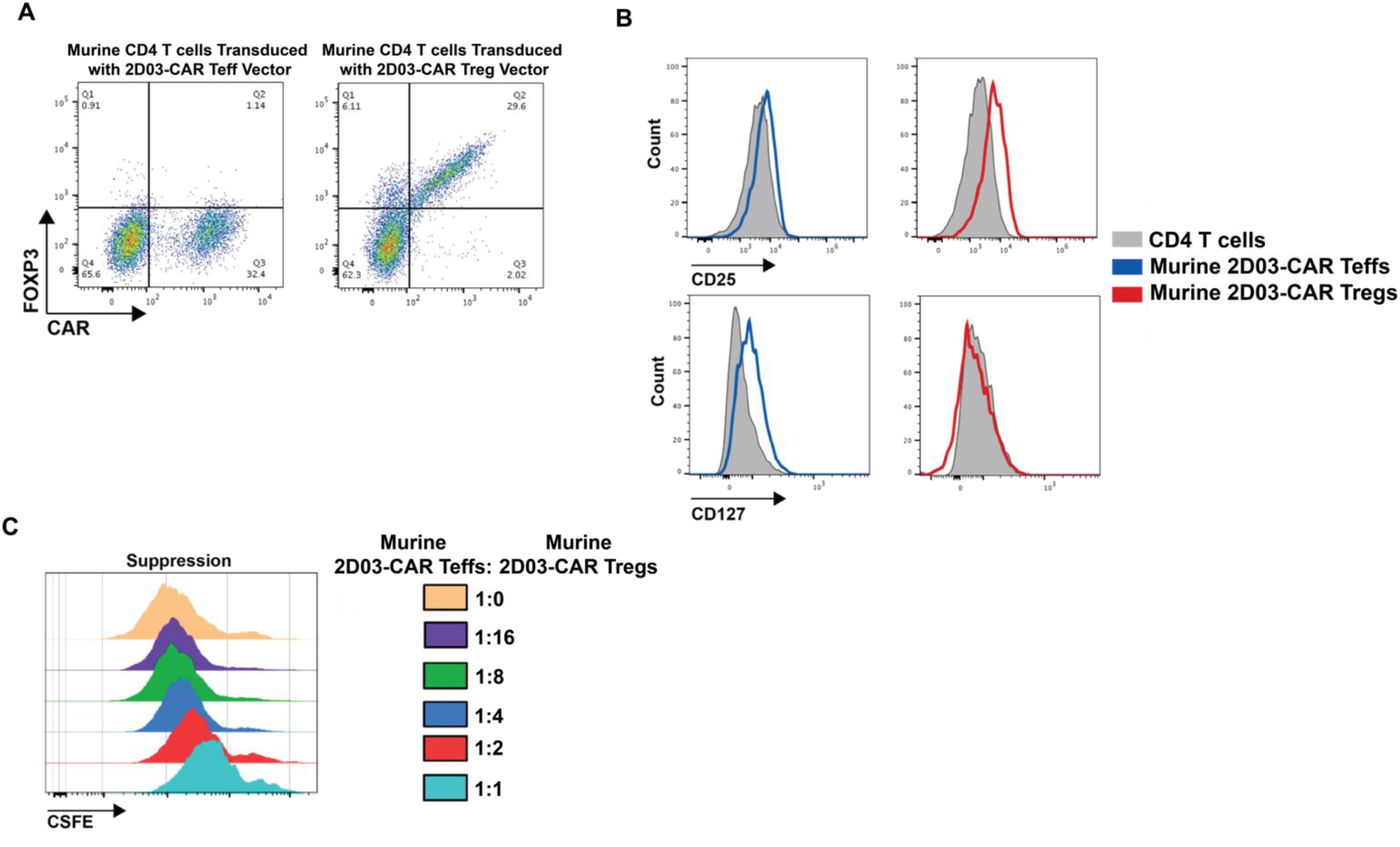
Murine 2D03-CAR Treg Development and Characterization. **A)** Representative flow plots of FOXP3 and CAR expression of transduced murine CD4 T cells. **B)** Murine 2D03-CAR Tregs were CD25^hi^ and CD127^low^ compared to murine CD4 T cells or murine 2D03-CAR Teffs **C)** CSFE-labeled murine 2D03-CAR Teffs were stimulated by MDA-ApoB100 and co-cultured with decreasing frequencies of murine 2D03-CAR Tregs.

**Supplementary Figure 2:**
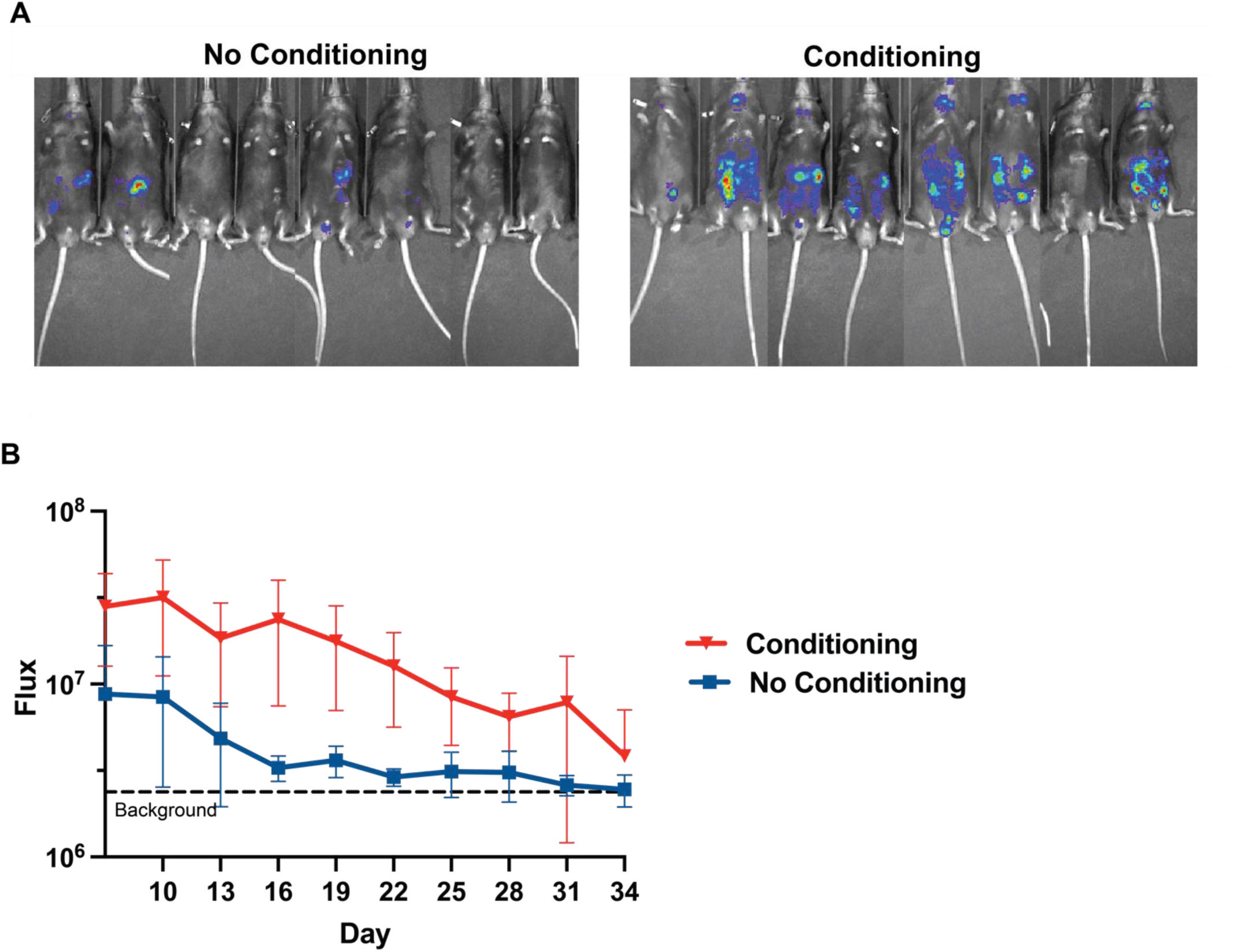
Murine 2D03-CAR Treg Persistence. **A)** Representative images of bioluminescence in conditioned or non-conditioned *C57BL/6J* mice one week after injection of T cells. **B)** Total flux corresponding to T cell persistence over 6 weeks of conditioned vs. non-conditioned mice.

## Acknowledgements

The authors want to acknowledge funding support from the Department of Veterans’ Affairs (I01 BX006247) (A.D.P.), Hematology Research Training Program T32 HL07439 (R.D.S.) and the American Society of Hematology Research Training Award for Fellows (R.D.S.). Additionally, we are grateful for editing contributions from Saar Gill, MD.

## Author contributions

R.D.S. and A.D.P. designed the research and wrote the manuscript. R.D.S., S-J.A.H., and X.B. performed the experiments. D.A.D., A.F., S.K.B., J.T.K., and F.L. provided experimental support. K.M. and D.J.R. advised the project and edited the manuscript.

